# Development of a replication-defective MPXV platform for fundamental and therapeutic research

**DOI:** 10.1101/2025.06.21.660854

**Authors:** Jiannan Chen, Liyuan Hu, Ning Shi, Jiaxin Tian, Yuyi Zhang, Sicheng Tian, Xinyu Cao, Zhuo Ha, Qiliang Cai, Yongxu Lu, Geoffrey L. Smith, Youhua Xie, Huijun Lu, Ping Zhang, Rong Zhang

## Abstract

The recent global outbreaks of mpox highlight the urgent need for both fundamental research and antiviral development. However, studying monkeypox virus (MPXV), with its large and complex genome, remains challenging due to the requirement for high-containment facilities. Here, we describe a novel strategy for *de novo* assembly of MPXV clade IIb genomes in bacterial artificial chromosomes using transformation-associated recombination cloning. Leveraging CRISPR-Cas9 and Lambda Red recombination, we engineered replication-defective MPXV particles with dual deletions of *OPG96* (*M2R*) and *OPG158* (*A32.5L*)—genes essential for virion assembly, that are capable of recapitulating key stages of the viral life cycle. Our work demonstrates the utility of replication-defective MPXV particles as a reliable platform for high-throughput antiviral discovery, offering significant advantages for both fundamental virology studies and therapeutic development against orthopoxviruses.

## INTRODUCTION

Monkeypox virus (MPXV), the causative agent of the zoonotic disease mpox, is an enveloped double-stranded DNA virus belonging to the *Poxviridae*, which also includes variola virus (VARV), vaccinia virus (VACV), and cowpox virus (CPXV) ^1^. MPXV strains are classified into two distinct clades: Clade I, found mainly in Central Africa (particularly the Democratic Republic of the Congo, DRC), is associated with severe clinical symptoms and substantial mortality (4–11%); Clade II, prevalent in West Africa, is characterized by milder symptoms and a mortality rate of less than 1% ^2, 3^. For over 50 years, MPXV outbreaks in Africa have been driven predominantly by zoonotic spillover and caused isolated, self-limiting outbreaks with poor human-to-human transmission ^4^.

In 2022, MPXV clade IIb caused the first widespread community transmission of mpox outside Africa ^5, 6^, prompting the World Health Organization (WHO) to declare mpox a Public Health Emergency of International Concern (PHEIC). Subsequently, due to the rising number of cases caused by clade Ib, the WHO declared a second mpox PHEIC in 2024 ^7^, following its earlier reprioritization of mpox as a neglected tropical disease with pandemic potential. This underscores the urgent need for effective prevention and treatment strategies ^8^.

However, the large and complex MPXV genome (∼200 kb), and the high biosafety containment (e.g., BSL-3) required to handle the live virus hinders studies on its molecular biology, pathogenesis, and antiviral drug development. Consequently, the development of tools such as replication-defective viruses—which enable genome manipulation and recapitulate viral infection under lower biosafety conditions— provides a valuable platform for scientific research.

Despite the global spread of MPXV, few drugs are approved for clinic use. Currently, only two therapeutics, tecovirimat (ST-246, TOPXX) and cidofovir or its prodrug brincidofovir (CMX001) are authorized for mpox treatment ^9^. Tecovirimat dimerizes the orthopoxvirus protein P37 (also called F13) encoded by the *F13L* gene (now renamed orthopoxvirus gene 57 (*OPG57*)^10^, to inhibit the formation of extracellular enveloped virions (EEVs) and cell-cell spread^11, 12^, but its efficacy remains unproven in clinical settings ^9, 13, 14^. Cidofovir or brincidofovir can inhibit viral DNA polymerase activity^15^, but is associated with gastrointestinal and hepatic toxicity. Thus, the identification of novel antivirals against MPXV is important.

To establish a safe and reliable alternative to authentic MPXV for antiviral research, we assembled a circular, full-length MPXV genome within a bacterial artificial chromosome (BAC) by transformation-associated recombination (TAR) cloning in yeast. We deleted MPXV homologs of *OPG96* (VACV *L2R*) and *OPG158* (VACV *A30.5L*) — both essential for viral envelope formation ^16, 17, 18^—to generate replication-defective MPXV (rdMPXV) particles that mimic the viral life cycle.

## RESULTS

### Assembly of the MPXV genome lacking *OPG96* (MPXV gene *M2R*)

Compared to clade I MPXV strains, MPXV clade IIb was associated with mild disease symptoms during the 2022 outbreak and so was selected for construction of rdMPXV particles. The entire viral genome was divided into 23 fragments (F1–F23), with 80–100 bp overlaps between adjacent fragments. The MPXV *OPG96* (*M2R*) lies within fragment F10, and encodes a 92 aa protein that is highly conserved in orthopoxviruses and essential for viral envelope formation and subsequent morphogenesis of IMV (intracellular mature virion) and EEV (extracellular enveloped virion) ^16, 19^. Therefore, this gene was selected for deletion from the MPXV genome. As illustrated in Fig. 1, adjacent fragments (grouped in sets of four), excluding F1, F2, and F11, were assembled by transformation-associated recombination (TAR) cloning in yeast to generate fragments B, C, E, F, and G. The reporter expression cassette, mGreenLantern (mGreen)-P2A-Gaussia luciferase, under the control of the viral late promoter P11, was inserted together with the linearized yeast/*E. coli* shuttle plasmid pBAC, into the *thymidine kinase* (*tk*) gene (*OPG101*) within fragment F11, yielding fragment D. Fragments B to G were then assembled by TAR cloning to generate intermediate plasmid pBAC-I^Δ96^, which lacks *OPG96* (Fig. 1).

**Figure 1.**
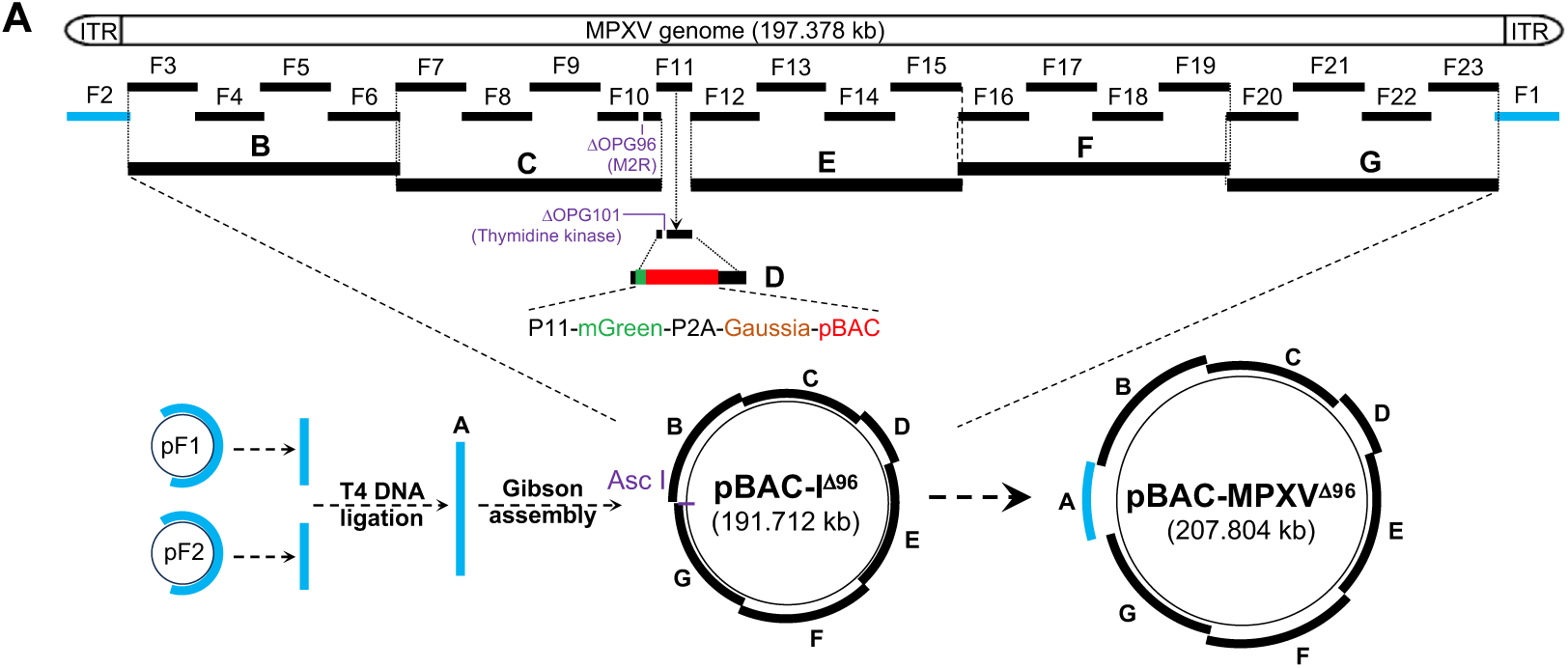
Assembly of the MPXV genome lacking *OPG96*(*M2R*). Schematic of the assembly of the MPXV genome. The full-length genome was divided into 23 fragments (F1–F23), with deletion of *OPG96* in F10. Every four adjacent fragments (except F1, F2, and F11) were assembled via TAR cloning in yeast to generate fragments B, C, E, F, and G. The mGreenLantern (mGreen)-P2A-Gaussia luciferase reporter cassette and the yeast/*E. coli* shuttle plasmid pBAC were inserted into the *tk* gene (*OPG101*) within F11, forming fragment D. Fragments B to G were further assembled through TAR cloning to obtain plasmid pBAC-I^Δ96^. Fragments F1 and F2 in plasmids were digested with SapI, ligated to form fragment A, and assembled with AscI-linearized pBAC-I^Δ96^ via Gibson assembly, yielding plasmid pBAC-MPXV^Δ96^, which contains the full-length MPXV genome with *OPG96* deleted.

The plasmid pBAC-I^Δ96^ exhibited stable maintenance in *E. coli* over 10 passages (Supplementary Fig. 1A), as confirmed by next-generation sequencing (NGS). No mutations or indels (insertions or deletions) were found in the viral genome or reporter region compared to the reference sequence (Supplementary Fig. 1B and 1C).

The remaining fragments F1 and F2 in plasmids were digested with the Type IIS restriction enzyme *SapI* and ligated *in vitro* to form fragment A. Fragment A was then inserted into the *AscI*-linearized pBAC-I^Δ96^, yielding the plasmid pBAC-MPXV^Δ96^, which contains the full-length MPXV genome lacking *OPG96* (Fig. 1).

### Generation of replication-defective MPXV particles lacking *OPG96*

To rescue the replication-defective virus particles, wild type CV-1 cells (CV-1-WT) were transduced with a lentivirus expressing OPG96 with codon-optimized coding sequences. The resulting stable cells (CV-1-96) were infected with fowlpox virus (FPV) to provide poxvirus transcriptional enzymes and then transfected with pBAC-MPXV^Δ96^ (Fig. 2A). The green fluorescence from the expression of mGreen reporter and cytopathic effect were monitored daily. On day 5 post-transfection, distinct viral plaques were observed (Fig. 2A), and the replication-defective MPXV particles lacking *OPG96* (rdMPXV^Δ96^) were harvested.

**Figure 2.**
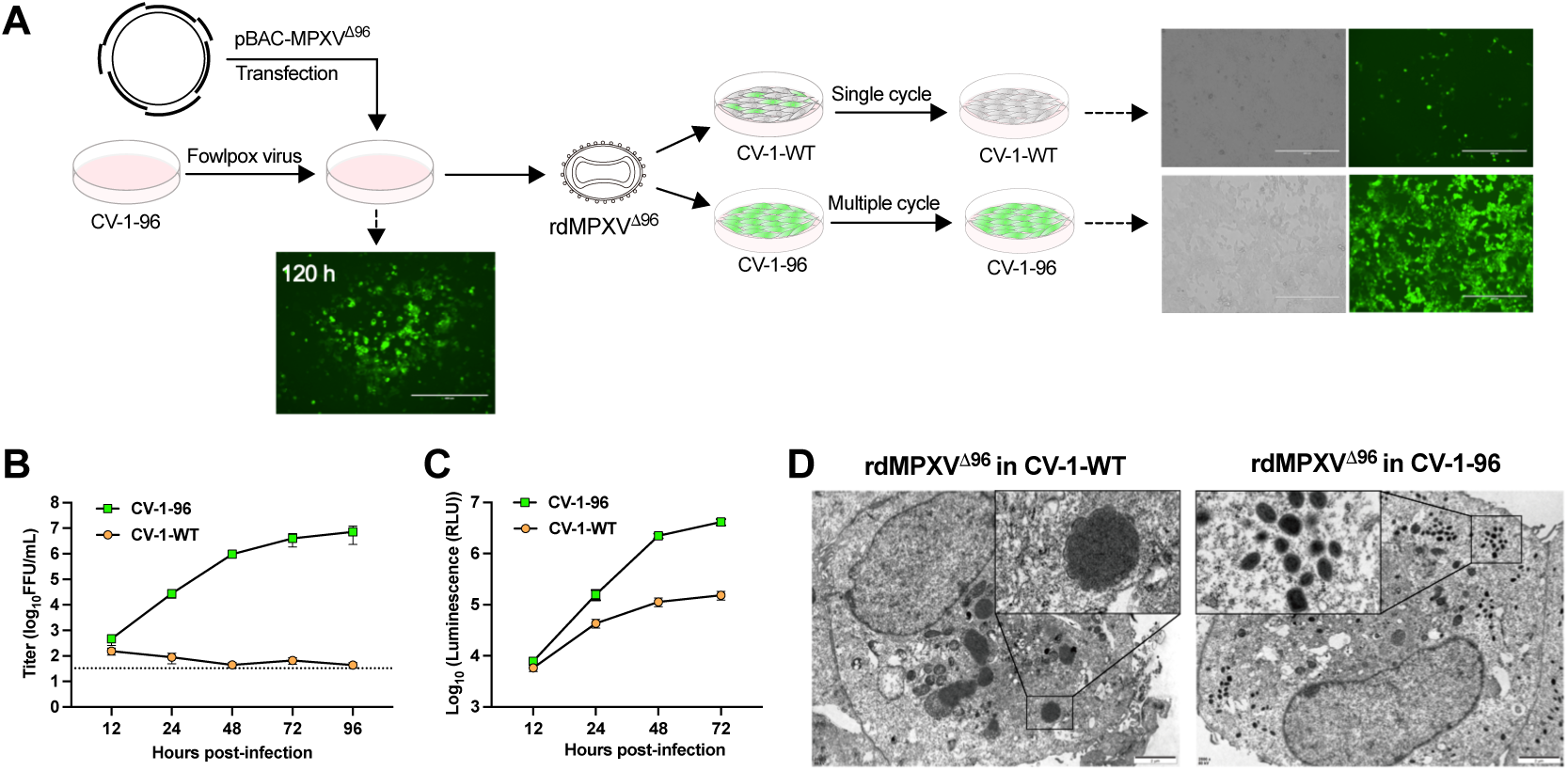
Generation of replication-defective MPXV particles lacking OPG96. **A.** Schematic of the generation of replication-defective MPXV particles (rdMPXV^Δ96^). CV-1-96 cells were infected with FPV for 2 h, followed by transfection with pBAC-MPXV^Δ96^. Cytopathic effects were observed at 120 h post-transfection, and rdMPXV^Δ96^ particles were harvested and used to infect CV-1-96 and CV-1-WT cells (MOI 0.05, 48 h). Representative fluorescence images were shown. Scale bar, 400 μm. **B-C.** Growth kinetics and Gaussia luciferase activity of rdMPXV^Δ96^ in CV-1-WT and CV-1-96 cells (MOI 0.05). Cells and supernatants were collected at the indicated time points for viral titration; supernatants alone were used for measuring luciferase activity. **D.** Transmission electron microscopy (TEM) of rdMPXV^Δ96^-infected CV-1-WT and CV-1-96 cells (MOI 1, 36 h). Representative images from two independent experiments are shown. Scale bar, 2 μm.

With the deletion of *OPG96*, which encodes a protein crucial for virion assembly, the rdMPXV^Δ96^ particles could only enter and express virus proteins in CV-1-WT cells, but were unable to form progeny IMV or EEV to infect neighboring cells. As expected, only single mGreen-positive cells were detected in CV-1-WT when compared to CV-1-96 cells which expressed OPG96 (Fig. 2A). The growth kinetics and Gaussia luciferase expression of rdMPXV^Δ96^ in CV-1-WT and CV-1-96 cells were measured. Abortive replication in CV-1-WT cells was observed, while the virus titer reached around 10^7^ FFU in CV-1-96 cells (Fig. 2B). Although moderate increase of luciferase activity was detected in CV-1-WT cells, the significant increase in CV-1-96 cells suggested that rdMPXV^Δ96^ underwent multiple cycles of replication, expressing abundant luciferase (Fig. 2C). Transmission electron microscopy was used to study the assembly of virions. Structurally intact virions were observed in CV-1-96 cells infected with rdMPXV^Δ96^ (Fig. 2D). For rdMPXV^Δ96^ infection in CV-1-WT cells, only short’spicule’-coated membrane arcs resembling crescent segments were detected around the large dense aggregates (Fig. 2D), which was consistent with the previous report describing a VACV mutant not expressing *OPG96* (VACV gene *L2R)* ^16^. Thus, these results demonstrated that rdMPXV^Δ96^ can only complete the entire lifecycle and propagate in OPG96 trans-complementing cells.

The defective replication property of rdMPXV^Δ96^ upon serial passaging was further confirmed in passage 1 (P1), P5, and P10 viruses, in which no viral amplification was detected in CV-1-WT cells (Supplementary Fig. 2A and 2B). PCR analysis of viral genomic DNA from P5 and P10 viruses revealed no indels in either the OPG96-deleted region or the reporter gene region (Supplementary Fig. 2C). Depth-of-coverage analysis of NGS reads also revealed no mutations or indels in the deleted OPG96 region and the whole genome compared to the rdMPXV^Δ96^ reference sequences (Supplementary Fig. 2D and 2E). Additionally, analysis of NGS reads indicated no introduction of FPV DNA into the P5 rdMPXV^Δ96^ genome (Supplementary Fig. 2F). The above results indicate that the replication-defective rdMPXV^Δ96^ particles exhibit stability in its trans-complementing cells.

### Generation of replication-defective MPXV particles with dual deletions of *OPG96* and *OPG158* (MPXV gene *A32.5L*)

The VACV protein *OPG158* (*A30.5*) interacts with *OPG96* (*L2*), and the deletion of *OPG158* also blocks the formation of the virion envelope, resulting in the failure to produce infectious viral particles ^17, 18^. To further strengthen safety, we constructed replication-defective virus particles lacking both *OPG96* and *OPG158* (MPXV gene *A32.5L*). The CRISPR/Cas9 gene editing in combination with Lambda Red recombination system were employed in *E. coli* to achieve this goal ^20^.

As illustrated in Figure 3, starting from *E. coli* DH10B containing the plasmid pBAC-I^Δ96^, the plasmid pEcCas, which constitutively expresses Cas9 and inducibly expresses the Lambda Red system, was transformed into DH10B for genome editing and recombination. The sgRNA-expressing plasmid pEcgRNA (targeting *OPG158*) and a homologous repair DNA template containing only the flanking sequences of the *OPG158* open reading frame (ORF) were then introduced into the cells. Cas9/sgRNA-mediated cleavage of *OPG158* within pBAC-I^Δ96^, combined with Lambda Red-induced homology-directed repair, resulted in deletion of *OPG158*. After removing both pEcgRNA and pEcCas plasmids in *E. coli*, the plasmid with dual deletions of *OPG96* and *OPG158* was obtained, validated by NGS (Supplementary Fig. 3A-C), and designated as pBAC-I^Δ96,158^. Using the virus rescue method applied for rdMPXV^Δ96^, replication-defective MPXV with double deletions (rdMPXV^Δ96,158^) was rescued in CV-1 cells trans-complementing both codon-optimized MPXV *OPG96* and *OPG158* (CV-1-96,158).

**Figure 3.**
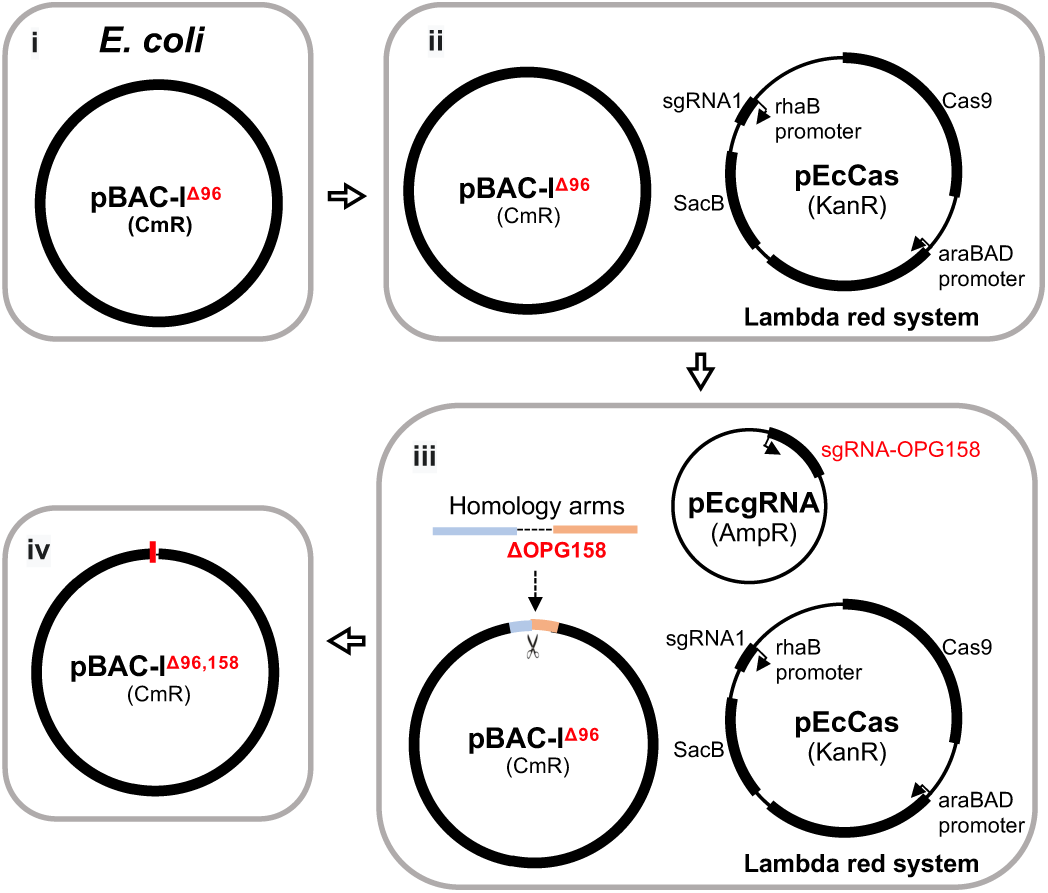
**CRISPR-based editing in combination with Lambda Red system to delete *OPG158* in plasmid pBAC-I^Δ96^ to construct pBAC-I**^Δ96,158^. (i-ii) DH10B cells were transformed with pBAC-I^Δ96^ and pEcCas (expressing Cas9 and Lambda Red recombination system); (iii) A homologous template (designed to delete *OPG158*) and pEcgRNA (expressing sgRNA targeting *OPG158*) were electroporated into DH10B cells. The Cas9/sgRNA complex cleaved *OPG158* in pBAC-I^Δ96^, and Lambda Red-mediated repair generated pBAC-I^Δ96,158^ with *OPG158* deleted; (iv) pEcgRNA and pEcCas were eliminated, yielding DH10B cells containing only pBAC-I^Δ96,158^.

Due to the dual-gene deletion, the rdMPXV^Δ96,158^ particles could theoretically only enter and express proteins in CV-1-WT and CV-1-96 cells, but be unable to form progeny virions unless propagated in the trans-complementing CV-1-96,158 cell line. This was confirmed by viral growth kinetics and Gaussia luciferase expression, which demonstrated that rdMPXV^Δ96,158^ could complete multi-cycle propagation in the CV-1-96,158 cells, but not in the CV-1-WT or CV-1-96 cells (Fig. 4A and 4B). Additionally, the defective replication of rdMPXV^Δ96,158^ was evaluated in dormice, a natural host for MPXV infection^21, 22^. Over 14 days, comparable increases in bodyweight were observed in mock-and rdMPXV^Δ96,158^-inoculated mice, whereas a significant reduction in body weight was noted in mice challenged with wild type MPXV (Fig. 4D). Furthermore, rdMPXV^Δ96,158^ did not exhibit productive replication in the lungs of dormice by day 14, in contrast to wild-type MPXV (Fig. 4C).

**Figure 4.**
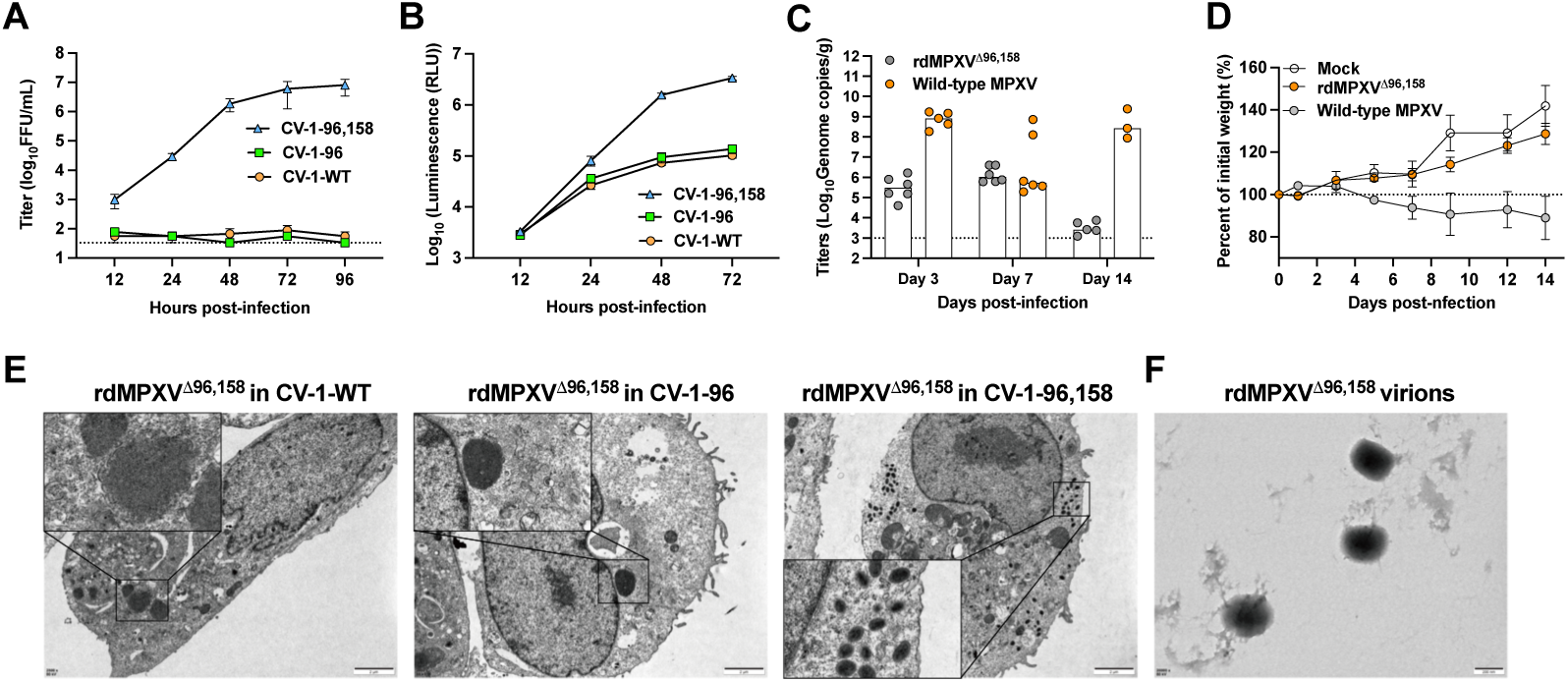
**Characterization of replication-defective MPXV particles with dual deletions of *OPG96* (*M2R*) and *OPG158* (*A32.5L*). A-B**. Growth kinetics (A) and Gaussia luciferase activity (B) of rdMPXV^Δ96,158^ in CV-1-WT, CV-1-96, and CV-1-96,158 cells (MOI 0.05). Cells and supernatants were collected for viral titration; supernatants were used for measuring luciferase activity. **C-D**. Viral replication in the lungs and body weight change of dormice. Dormice aged 10-12 weeks were inoculated intranasally with PBS (mock), 5×10^5^ focus-forming unit (FFU) of rdMPXV^Δ96,158^ particles, or 1×10^5^ FFU of wild type MPXV. Viral genomic copies in the lungs were determined by qPCR on day 3, 7, and 14 post-infection (C), and body weight was monitored over 14 days (D). **E-F**. Transmission electron microscopy (TEM) analysis of rdMPXV^Δ96,158^– infected cells (E) and virions (F). CV-1-WT, CV-1-96, and CV-1-96,158 cells were infected with rdMPXV^Δ96,158^ (MOI 1, 36 h) and fixed with 2.5% glutaraldehyde for TEM. rdMPXV^Δ96,158^ particles harvested from CV-1-96,158 cells were used for TEM. Representative images from two experiments are shown. Scale bar, 2 μm. Data in A and B are from three independent experiments, and each performed in triplicate.

Transmission electron microscopy was used to examine virion assembly. Structurally intact virions were observed in rdMPXV^Δ96,158^–infected CV-1-96,158 cells, but not in both CV-1-WT and CV-1-96 cells (Fig. 4E), consistent with previous reports on VACVs lacking either gene ^16, 18^. Additionally, intact virions released from rdMPXV^Δ96,158^–infected CV-1-96,158 cells were detected (Fig. 4F).

The stability of replication-defective rdMPXV^Δ96,158^ was studied. After 10 passages in CV-1-96,158 cells, rdMPXV^Δ96,158^ maintained the single-cycle replication property in CV-1-WT and CV-1-96 cells (Supplementary Fig. 3D). PCR analysis of the genomic DNA from the P10 virus confirmed no indels in *OPG96*, *OPG158*, and reporter regions (Supplementary Fig. 3E). NGS analysis further revealed no mutations and indels in the deleted *OPG96* and *OPG158* regions, and the whole genome of the P5 and P10 rdMPXV^Δ96,158^ compared to the rdMPXV^Δ96,158^ reference sequences (Supplementary Fig. 3F-H). Thus, these results demonstrate that rdMPXV^Δ96,158^ exhibits favorable safety and stability. Because of dual-gene deletions, rdMPXV^Δ96,158^ possesses a higher safety profile compared to rdMPXV^Δ96^, which has only a single gene deleted.

### Application of rdMPXV^Δ96,158^ particles in cell biology and antiviral assessment

To expand the application of replication-defective rdMPXV^Δ96,158^ particles, the susceptibility of commonly used cell lines to rdMPXV^Δ96,158^ infection was evaluated. As shown in Figure 5A and Supplementary Figure 4A, HaCaT, CV-1, Vero E6, BS-C-1, and HFF-1 cells were highly susceptible to infection, as indicated by expression of mGreen reporter, while 293T, Huh7, and SW13 cells exhibited low susceptibility. It is intriguing that the susceptibility of skin and fibroblast cells such as HaCaT and HFF-1 may correlate with the MPXV tissue tropism. Previous studies have reported that heparan sulfate and chondroitin sulfate are involved in the attachment of VACV virions to the cell membrane ^23, 24, 25, 26^. The *B3GAT3* gene, responsible for heparan sulfate and chondroitin sulfate biosynthesis ^27^, was found to be important for MPXV infection, as its deletion reduced the infection efficiency of rdMPXV^Δ96,158^ in A549 cells (Fig. 5B and Supplementary Fig. 4B).

**Figure 5.**
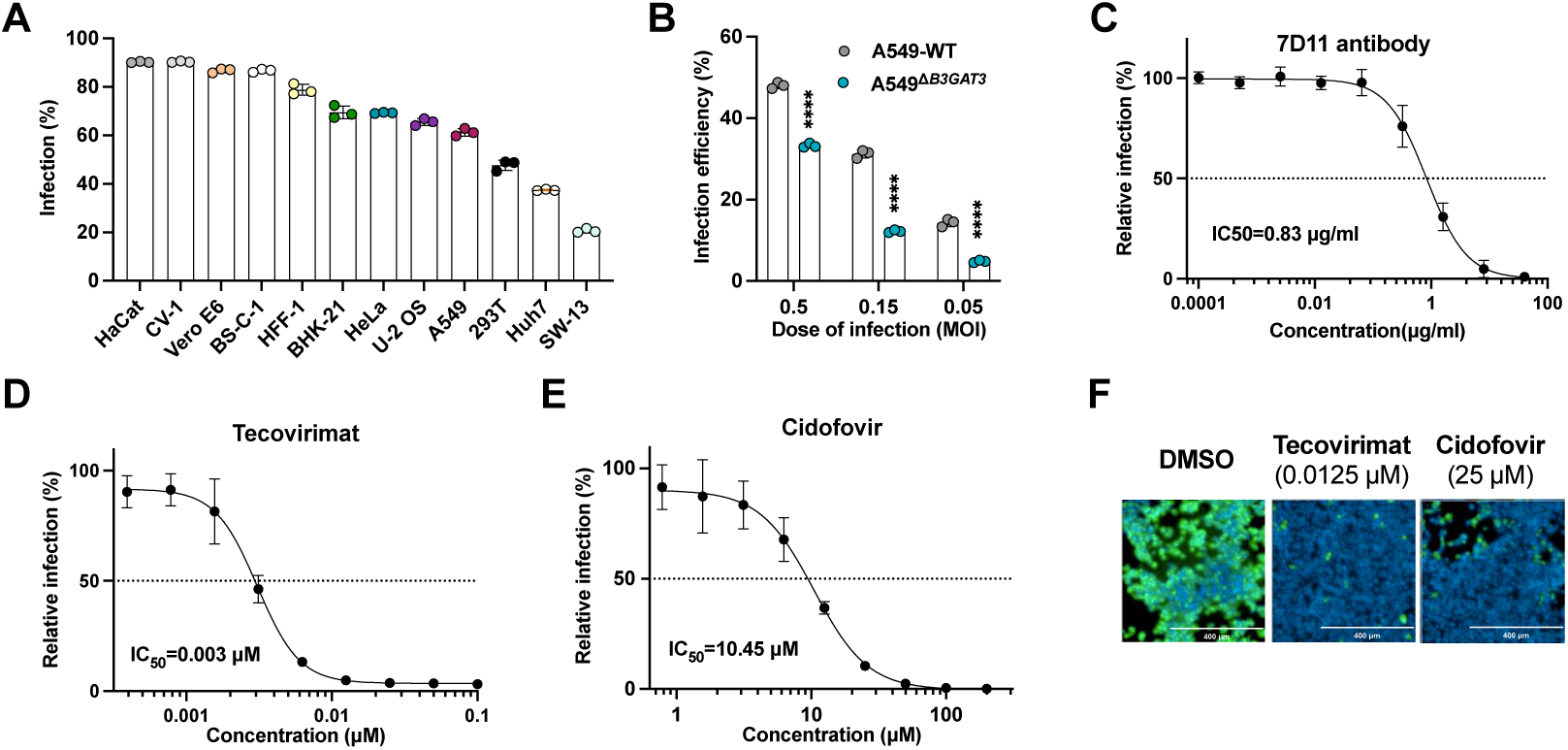
Application of rdMPXV^Δ96,158^ particles in cell biology and antiviral assessment. **A**. Susceptibility of cells to rdMPXV^Δ96,158^ (MOI 0.5, 48 h) analyzed by flow cytometry. B. Infection efficiency of rdMPXV^Δ96,158^ in WT and Δ*B3GAT3* A549 cells (MOI 0.5, 0.15, 0.05; 48 h). **C-E**. IC50 of 7D11 antibody (C), tecovirimat (D), and cidofovir (E) in CV-1-96,158 cells infected with rdMPXV^Δ96,158^ (MOI 0.01 or 0.1, 30 h). **F**. Fluorescence microscopy of CV-1-96,158 cells pre-treated with tecovirimat (0.125 μM) or cidofovir (25 μM), then infected with rdMPXV^Δ96,158^ (MOI 0.1, 30 h). Cells were fixed and stained with DAPI. Scale bar, 400 μm. Data in A-E are from three independent experiments, and each performed in triplicate. In panel B, significance was determined by unpaired t-test (n=3; ****P < 0.0001; mean ± s.d.).

Next, the capacity of rdMPXV^Δ96,158^ particles for antiviral development was explored. In VACV, the specific neutralizing antibody 7D11 targeting the L1 viral protein (*OPG95*) can block infection ^28^. This antibody also inhibited rdMPXV^Δ96,158^ infection with an IC_50_ of 0.83 μg/mL (Fig. 5C). Tecovirimat, which targets the orthopoxvirus P37 protein encoded by OPG57 (VACV *F13L*) and blocks wrapping of IMV, so that the intracellular enveloped virus (IEV, also called wrapped virus, WV) is not formed, and cidofovir, which targets viral DNA polymerase to inhibit DNA synthesis, are FDA-approved drugs for emergency use in MPXV patients ^9, 29^. Both drugs effectively inhibited rdMPXV^Δ96,158^ spread (tecovirimat) or replication (cidofovir), with IC_50_ values of 0.003 μM (tecovirimat) and 10.450 μM (cidofovir) in CV-1-96,158 cells (Fig. 5D-F).

Similar results were observed in human keratinocytes (HaCaT) stably-expressing OPG96 and OPG158 (HaCaT-96,158) (Supplementary Fig. 4C-E). The antiviral activities of tecovirimat and cidofovir assessed using replication-defective rdMPXV^Δ96,158^ particles were consistent with previous reports using wild-type MPXV in HFF and HFK cells ^29^.

## DISCUSSION

In this study, the full-length MPXV genome was assembled in a BAC plasmid with deletions of both OPG96 and OPG158, and used to generate replication-defective rdMPXV^Δ96,158^ particles that undergo normal replication in trans-complementing cells. The single-cycle replication property of rdMPXV^Δ96,158^ with dual gene deletions enables its use in lower-biosafety-level facilities (e.g., BSL-2).

VACV, one of the most extensively studied poxviruses, provides a foundational model for constructing replication-defective viral particles. Genes that are essential for viral replication include: a. genes involved in DNA replication, such as *OPG116* (*D4*) ^30, 31^, *OPG110* (*H5*) ^32^, and *OPG79* (*I3*) ^33^; b. genes related to intermediate-stage viral gene transcription, such as *OPG134* (*A8*) and *OPG150* (*A23*) ^34^; and c. genes involved in virion morphogenesis, such as *OPG96* (*L2*) ^16, 18^, *OPG158* (*A30.5*) ^17, 18^, *OPG112* (*H7*) ^35^, and *OPG78* (*I2*) ^36^. To facilitate the subsequent use of replication-defective particles to investigate the life cycle of MPXV, we selected *OPG96* and *OPG158* for trans-complementation. These genes are distantly located within the MPXV genome and are both required for virion morphogenesis. To minimize the risk of generating replication-competent virus through recombination, we codon-optimized the deleted genes and expressed them via separate lentiviral vectors in cells for trans-complementation.

Most existing replication-defective VACVs were derived from infectious virus, where the viral genome was edited within complementing cell lines, with replication-defective particles obtained after multiple purification rounds ^16, 18, 36^. Additionally, rescue of live poxviruses could be achieved in helper virus-infected cells by transfecting either segmented genomic DNA with homologous arms ^37^ or circularized genomes containing BAC elements derived from concatemeric viral DNA ^38^. In contrast, we employed yeast TAR cloning to assemble the full-length MPXV genome *de novo*. Due to the presence of inverted terminal repeats (ITRs) in fragments F1 and F2, we first assembled the genome without these fragments into a circular intermediate plasmid (pBAC-I^Δ96^), which was stable upon successive passaging in *E. coli.* The remaining F1 and F2 fragments were then inserted to construct pBAC-MPXV^Δ96^, which was used to produce replication-defective particles.

Traditional methods for constructing mutant poxviruses rely on homologous recombination in infected cells using selection markers ^39^, an approach limited by low efficiency, the need for multiple purification steps for generating mutants. Similarly, conventional editing of BAC plasmids bearing VACV genomes depends on homologous recombination with selection markers ^40^, restricting iterative modifications. To overcome these limitations, we developed a platform combining CRISPR-based genome editing with Lambda Red recombination system in *E. coli*, enabling rapid, efficient, and multiplexed modifications of the stable intermediate plasmid pBAC-I^Δ96^. This system significantly facilitates manipulation of the MPXV genome, and is applicable to other BAC systems containing large viral genomes, such as herpesviruses and coronaviruses ^41, 42^.

Our engineered replication-defective MPXV particles establish a robust, convenient, and reliable platform for high-throughput antiviral discovery, offering significant advantages for both fundamental virology studies and therapeutic development against orthopoxviruses.

## METHODS

### Cells and viruses

CV-1, HaCaT, BS-C-1, SW13, Huh7, Vero E6 (Cell Bank of the Chinese Academy of Sciences, Shanghai, China), HFF-1 (ATCC #SCRC-1041), U-2 OS (ATCC #HTB-96), BHK-21 (ATCC #CCL-10), HEK 293T (ATCC #CRL-3216), A549 (ATCC #CCL-185), and HeLa (ATCC #CCL-2) were cultured at 37°C in Dulbecco’s Modified Eagle’s Medium (Hyclone) supplemented with 10% fetal bovine serum (FBS), 10 mM HEPES, 1 mM sodium pyruvate, 1× non-essential amino acids, and 100 U/mL penicillin-streptomycin. All cell lines were routinely tested and confirmed free of mycoplasma contamination. Fowlpox virus (FPV) was propagated in primary chick embryo fibroblasts. MPXV IIb strain (GenBank: PP778666.1) was grown in Vero E6 cells.

### Transformation-associated recombination (TAR) cloning of plasmids

The complete genome of MPXV clade IIb, with a length of 197,378 bp, was divided into 23 fragments (F1–F23). Adjacent fragments were designed with homologous arms, and *OPG96* (*M2R*) in fragment F10 was deleted. As illustrated in Figure 1, every four adjacent fragments (except F1, F2, and F11) were assembled in *Saccharomyces cerevisiae* strain VL6-48 using the yeast/*E. coli* shuttle plasmid pBAC. Assembled plasmids were purified from VL6-48 and transformed into *E. coli* DH10B for amplification, resulting in plasmids designated pBAC-B,-C,-E,-F, and-G. For pBAC-D construction, the reporter cassette (mGreenLantern-P2A-Gaussia luciferase) under the control of the late viral promoter P11 was synthesized and inserted into OPG101 (*J2R* gene, *thymidine kinase* (*tk*)) of fragment F11. The modified F11 fragment was cloned into the shuttle plasmid. Fragments B–G (digested from pBAC-B to pBAC-G) were co-transformed with linearized pBAC-D into yeast. Assembled plasmids were purified and transformed into DH10B, yielding the final plasmid pBAC-I^Δ96^, which lacks F1, F2, and *OPG96*.

### Yeast transformation

Yeast transformation via lithium acetate was performed according to Gietz, R. D. et al. ^43^. Briefly, VL6-48 cells were grown in 2× YPAD medium, washed, and resuspended in sterile water. A 100 μL yeast suspension was transferred to a microcentrifuge tube and mixed with transformation solution containing 36 μL lithium acetate (1.0 M), 240 μL PEG 3350 (50% w/v), 50 μL single-stranded carrier DNA (2.0 mg/mL), and 34 μL DNA (fragments and linearized shuttle vector). The mixture was heat-shocked at 42°C for 40 min to facilitate DNA uptake. Cells were harvested by centrifugation, resuspended in sterile water, plated on selective medium, and incubated at 30°C for 3–4 days. Positive clones were validated by PCR and sequencing. Plasmids were extracted and electroporated into *E. coli* DH10B for amplification.

### CRISPR-based editing of the plasmid in *E. coli*

To knock out *OPG158* (*A32.5L*) via homologous recombination, a DNA template encompassing only the upstream and downstream sequences of *OPG158* was PCR-amplified and joined by overlapping-extension PCR. The spectinomycin resistance gene in the pEcgRNA vector (Addgene #166581) was replaced with an ampicillin resistance gene, and an sgRNA (5’-TTTGGAACGTTTGATAAAAG-3’) targeting *OPG158* was cloned to generate pEcgRNA-sg*OPG158*. The plasmids pBAC-I^Δ96^ (constructed above) and pEcCas (Addgene #73227) were transformed into DH10B cells, and the Lambda Red recombination system was induced with 10 mM L-arabinose. The sgRNA-expressing plasmid pEcgRNA-sg*OPG158* and the DNA recombination repair template (lacking *OPG158* coding sequences) were co-electroporated into DH10B cells harboring pBAC-I^Δ96^ and pEcCas. Positive clones lacking *OPG158* were identified by PCR and sequencing. After overnight culture in selection medium, the pEcgRNA-sg*OPG158* and pEcCas plasmids were eliminated, resulting in DH10B cells containing only the plasmid pBAC-I^Δ96,158^ with dual deletions of *OPG96* and *OPG158*.

### Assembly of full-length MPXV genome and virus rescue

Fragments F1 and F2 in plasmids were digested with *SapI* and ligated using T4 DNA ligase to form fragment A. Fragment A and the *AscI*-linearized pBAC-I^Δ96^ plasmid were assembled via Gibson assembly to generate the plasmid pBAC-MPXV^Δ96^, which contains the full-length MPXV genome lacking OPG96. Similarly, the pBAC-MPXV^Δ96,158^ plasmid lacking *OPG96* and *OPG158* was constructed in the pBAC-I^Δ96,158^ backbone. CV-1-96 or CV-1-96,158 stable cell lines were infected with FPV for 2 h. After washing, the pBAC-MPXV^Δ96^ or pBAC-MPXV^Δ96,158^ plasmid was transfected into the cells, and the presence of green fluorescence and cytopathic effect (CPE) was monitored daily. Supernatants and cells were harvested and subjected to three freeze-thaw cycles at −80°C to generate the P0 virus stock. The resulting replication-defective viral particles—with a single deletion of OPG96 (rdMPXV^Δ96^) or dual deletions of *OPG96* and *OPG158* (rdMPXV^Δ96,158^)—were designated accordingly.

### Viral growth kinetics and luciferase assay

In 12-well plates, cells were infected with the virus at an MOI of 0.05 for 2 h, followed by a medium change to DMEM containing 2% FBS. At the indicated time points, cells and supernatants were collected together and subjected to three freeze-thaw cycles at −80°C to release the virus. To determine the titer, monolayers of CV-1-96 or CV-1-96,158 in 96-well plates were inoculated with serially diluted virus for 2 h and then overlaid with methylcellulose for 36 h. Cells were fixed with 4% paraformaldehyde (PFA) in PBS for 1 h and washed with PBS. Green fluorescent foci were counted. To measure Gaussia luciferase activity, only the supernatants were collected. The luciferase activity was determined using the Secrete-Pair Gaussia Luciferase Assay Kit (iGeneBio #LF061) according to the manufacturer’s instructions. Briefly, 10 μL of the supernatant was added to a 96-well plate, followed by 100 μL of assay reagent, which was gently pipetted to mix thoroughly. After incubation at room temperature for 1 min, luminescence was recorded using a FlexStation 3 (Molecular Devices) with an integration time of 1 second per well.

### Electron Microscopy

CV-1-WT and CV-1-96 cells were infected with rdMPXV^Δ96^ at an MOI of 1. CV-1-WT, CV-1-96, and CV-1-96,158 cells were infected with rdMPXV^Δ96,158^ at an MOI of 1. At 36 h post-infection, cells were scraped and fixed with 2.5% glutaraldehyde for 1 h. Samples were further fixed with 1% osmium tetroxide at 20°C for 2 h, dehydrated through a graded ethanol series, and infiltrated overnight with acetone. After polymerization at 60°C for 48 h, ultrathin sections were prepared using a Leica UC7 microtome. Sections were stained with uranyl acetate and lead citrate, dried, and imaged using a FEI Tecnai G2 20 Twin transmission electron microscope.

### Assessment of cell susceptibility

The cell lines from different species and organs were infected with rdMPXV^Δ96,158^ at an MOI of 0.5 for 48 h. Representative images were captured using a fluorescence microscope. To determine infection efficiency, cells were trypsinized, fixed with 2% paraformaldehyde (PFA) for 10 min, and analyzed by flow cytometry. To examine the role of heparan sulfate and chondroitin sulfate in MPXV infection, a *B3GAT3*-knockout clonal cell line was generated in A549 cells by introducing a ribonucleoprotein (RNP) complex containing Cas9 protein and crRNA. Following limiting dilution and expansion, clonal knockout cells were selected and validated by sequencing.

### Assessment of viral stability

Trans-complemented stable cells in 12-well plates were infected with replication-defective MPXV particles at an MOI of 0.01 for 2 h. After washing, cells were maintained in DMEM supplemented with 2% FBS to allow viral propagation for 96 h, followed by three freeze-thaw cycles at −80°C to release virions. Virus particles were then used to infect new cells at an MOI of 0.01 for passaging as described. Genomic DNA from passage 5 (P5) and passage 10 (P10) was extracted. Regions encompassing *OPG96, OPG158*, or the mGreenLantern-P2A-Gaussia luciferase reporter cassette were amplified for gel electrophoresis and Sanger sequencing. Genomic DNA was also subjected to Next-Generation Sequencing (NGS) to analyze mutations or contamination.

### Plaque reduction neutralization (PRNT) assay

The neutralizing antibody 7D11 targeting OPG95 (VACV L1) was serially diluted in DMEM containing 2% FBS, starting from 80 μg/ml in 5-fold increments. The diluted antibody was mixed in equal volumes with 2% FBS DMEM containing 240 focus-forming units (FFU) of rdMPXV^Δ96,158^ (final volume: 100 μL) and incubated at 37°C for 1 h. CV-1-96,158 cells in a 96-well plate were infected with the mixture for 2 h, overlaid with methylcellulose for 30 h, and fixed with 4% PFA. Plaques were quantified using an ImmunoSpot^®^ Analyzer.

### Virus infection in animals

To evaluate the replication of rdMPXV^Δ96,158^ *in vivo*, dormice aged 10-12 weeks were inoculated intranasally with PBS (mock control), 5×10^5^ focus-forming unit (FFU) of rdMPXV^Δ96,158^ particles, or 1×10^5^ FFU of wild-type MPXV IIb strain (GenBank: PP778666.1) in a volume of 30 μl. Body weight was monitored over 14 days. Lung tissues were harvested on day 3, 7, and 14 post-infection for DNA extraction. Viral genomic copies were quantified by qPCR targeting OPG47 (F3L) using the following primers and probe: Fwd, 5’-CTCATTGATTTTTCGCGGGATA-3’; Rev, 5’-GACGATACTCCTCCTCGTTGGT-3’; Probe, 5’-CATCAGAATCTGTAGGCCGT-3’. All animal experiments were conducted in accordance with the National Institutes of Health (NIH) Guide for the Care and Use of Laboratory Animals and were approved by the Animal Care and Use Committees of the Changchun Veterinary Research Institute.

### Statistical analysis

Statistical significance was assigned when P values were < 0.05 using Prism Version 9 (GraphPad). Data analysis was determined by an ANOVA or unpaired t-test depending on data distribution and the number of comparison groups.

## ACKNOWLEDGEMENTS

Grants from the Program of Shanghai Academic Research Leader (22XD1420600 to R.Z.), Shanghai Municipal Science and Technology Major Project (ZD2021CY001), and Non-profit Central Research Institute Fund of Chinese Academy of Medical Sciences (2023-PT310-02) supported this work. We wish to acknowledge Xiaoqing Sun at Key Laboratory of Medical Molecular Virology (MOE/NHC/CAMS), Shanghai Frontiers Science Center of Pathogenic Microorganisms and Infection, School of Basic Medical Sciences of Fudan University for the help with next generation sequencing and data analysis. We would like to thank Guangzhou Eighth People’s Hospital of Guangzhou Medical University for providing MPXV samples. We also thank colleagues at the Biosafety Level 3 Laboratory of Changchun Veterinary Research Institute of Chinese Academy of Agricultural Sciences for help with the experiments.

## AUTHOR CONTRIBUTIONS

J.C., L.H., N.S., J.T., X.C., Z.H., and Y.L. performed the experiments. J.C., L.H., R.Z. designed the experiments. Y.Z. and S.T. provided technical or material support. Y.L., G.L.S., Y.X., H.L., P.Z., and R.Z. provided administrative, supervision support. J.C, L.H., N.S., and R.Z. performed data analysis. J.C. and R.Z. wrote the initial draft of the manuscript, with the other authors contributing to editing into the final form.

## SUPPLEMENTARY FIGURES LEGENDS

**Supplementary Figure 1.**
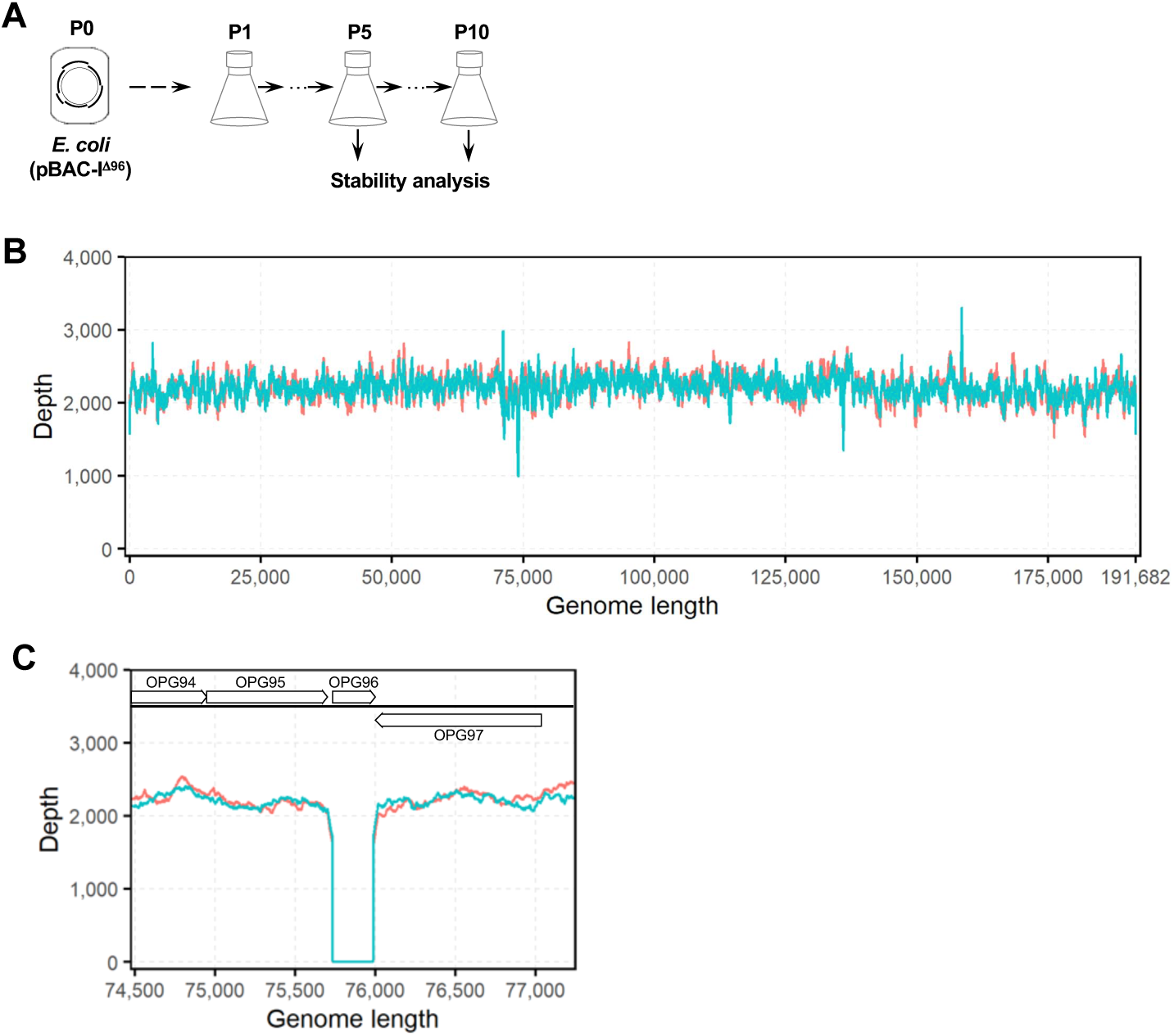
Stability analysis of the pBAC-IΔ96 plasmid. **A**. Schematic diagram of pBAC-I propagation in *E. coli* across generations. **B**. Read depth of *E. coli*-derived P5 (red line) and P10 (green line) of pBAC-I^Δ96^ aligned to the pBAC-I^Δ96^ reference sequences. **C**. Read depth of *E. coli*-derived P5 (red line) and P10 (green line) of pBAC-I^Δ96^ aligned to MPXV reference genome, with focus on OPG96 (gene M2R) and flanking regions. After filtering and pairing, the reads were aligned to the designed plasmid reference using the BWA software (version 0.7.18) with default parameters^44^.The coverage across different regions of the plasmid was then calculated using the bam-readcount tool ^45^, and the data were visualized using R.

**Supplementary Figure 2.**
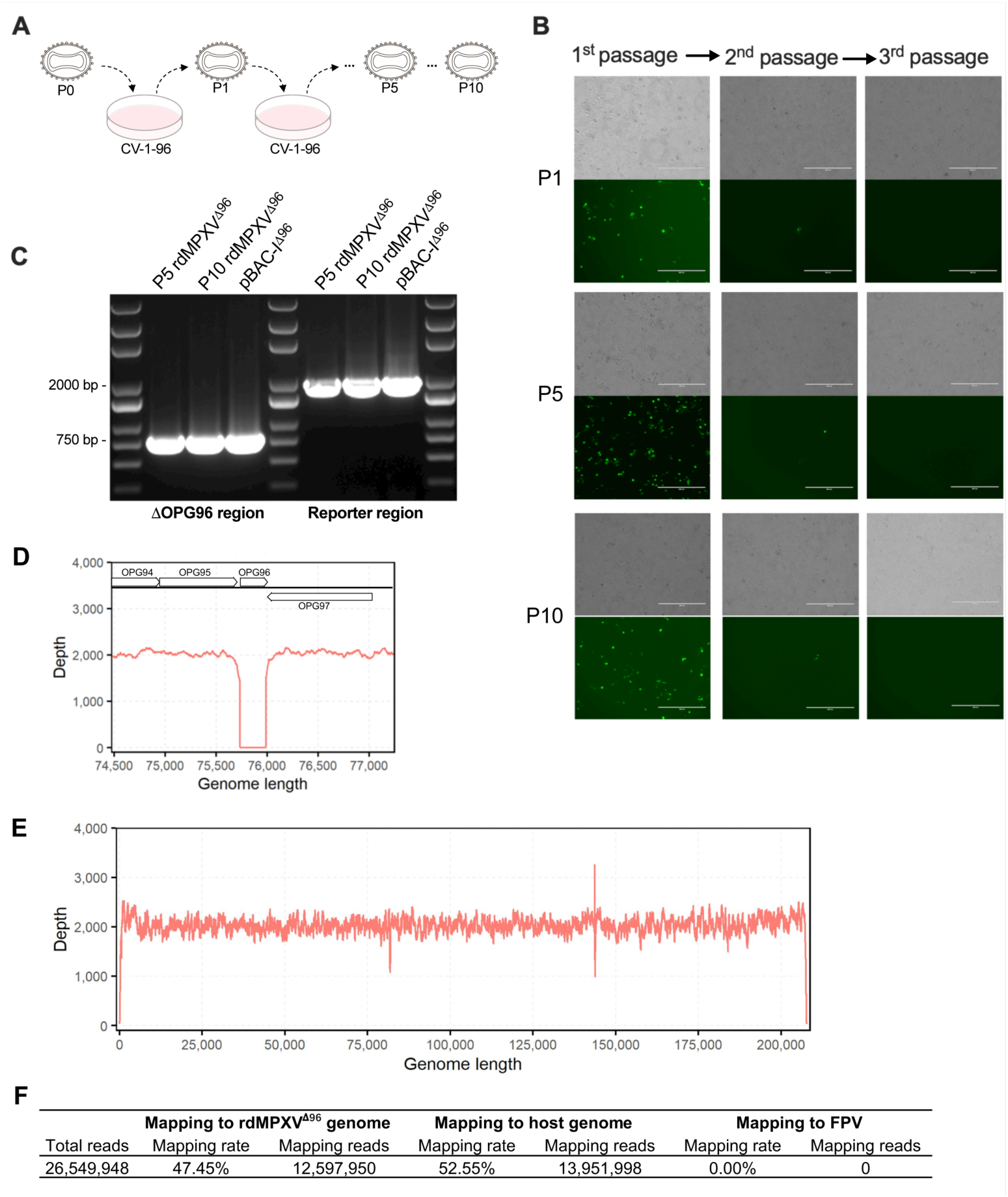
Stability of rdMPXVΔ96 upon serial passaging. **A.** Schematic of serial passaging of rdMPXV^Δ96^ in CV-1-96 cells. The P0 virus (harvested from CV-1-M cells transfected with pBAC-MPXV^Δ96^) was used to infect fresh CV-1-96 cells, yielding P1. Similarly, P5 and P10 viruses were obtained. **B.** Serial passaging of rdMPXV^Δ96^ in CV-1-WT cells. P1, P5, and P10 viruses from CV-1-96 cells (panel A) were passaged for three rounds in CV-1-WT cells. Brightfield and fluorescent images were captured at 96 h post-infection. Scale bar, 400 μm. **C.** PCR-based detection of the OPG96-deleted region and mGreenLantern-P2A-Gaussia luciferase reporter cassette in P5 and P10 rdMPXV^Δ96^. Viral genomic DNA from P5 and P10 was amplified by PCR, with pBAC-I^Δ96^ as a positive control. **D.** Read depth of P5 rdMPXV^Δ96^ aligned to MPXV reference genome, focusing on OPG96 and flanking regions. **E.** Read depth of P5 rdMPXV^Δ96^ aligned to rdMPXV^Δ96^ reference sequences. **F.** Analysis of sequencing reads from P5 of rdMPXV^Δ96^ mapping to virus or host genome.

**Supplementary Figure 3.**
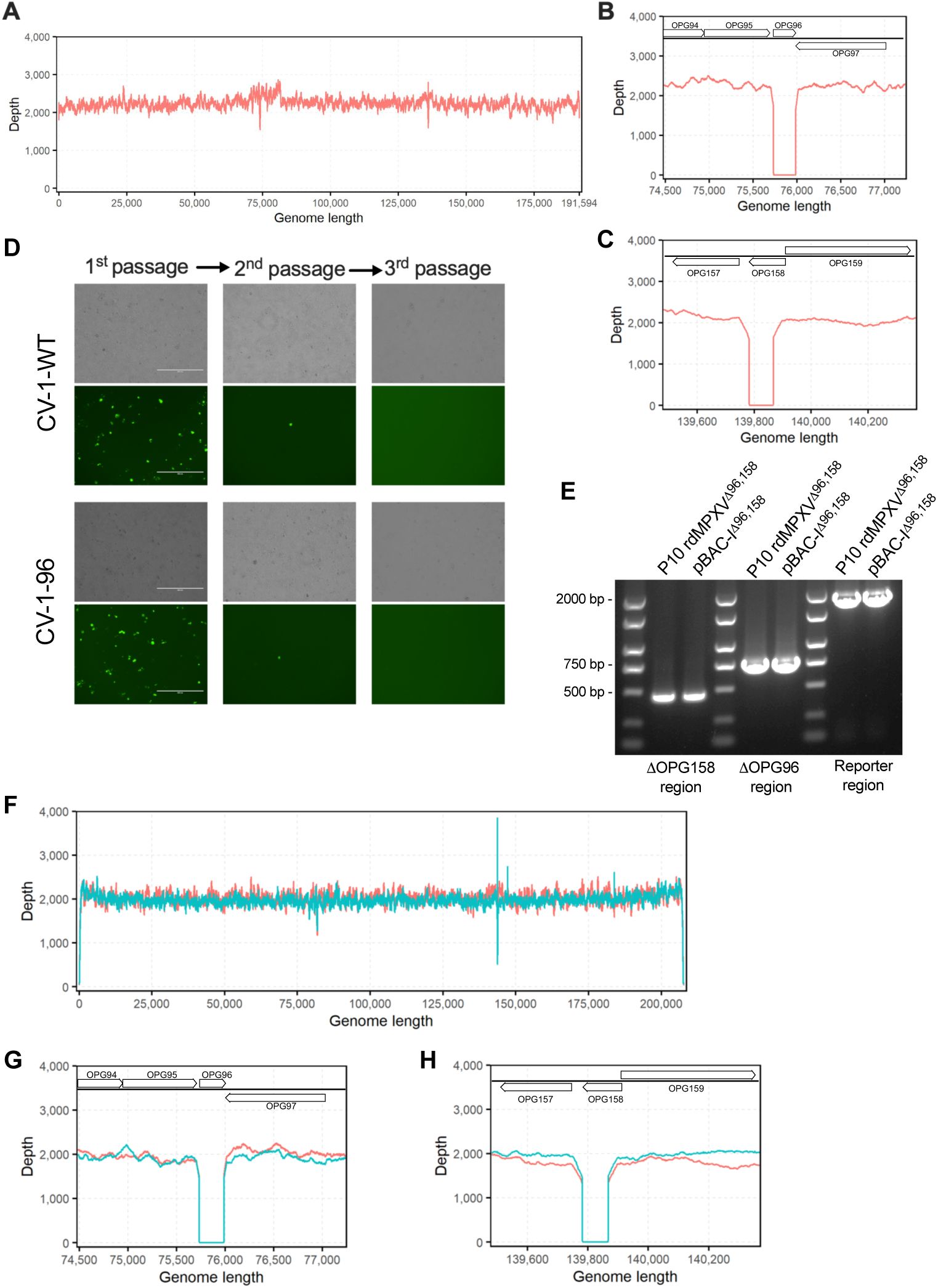
Sequence verification of pBAC-IΔ96,158 plasmid and stability analysis of rdMPXVΔ96,158 upon serial passaging. **A.** Read depth of *E. coli*-derived pBAC-I^Δ96,158^ aligned to pBAC-I^Δ96,158^ reference sequences. pBAC-I^Δ96,158^ plasmid was constructed by CRISPR-based editing in combination with Lambda Red recombination. **B-C.** Read depth of *E. coli*-derived pBAC-I^Δ96,158^ aligned to the MPXV reference genome, focusing on *OPG96* (B) and *OPG158* (gene A32.5L) (C) and their flanking regions. **D.** Stability analysis of serially passaged rdMPXV^Δ96,158^ in CV-1-WT and CV-1-96 cells. rdMPXV^Δ96,158^ passaged 10 times (P10) in CV-1-96,158 cells, was used to passage another three rounds in CV-1-WT and CV-1-96 cells. Brightfield and fluorescent images were captured at 96 h post-infection. Scale bar, 400 μm. **E.** PCR verification of *OPG96* and *OPG158* deletions and the mGreenLantern-P2A-Gaussia luciferase reporter cassette in P10 rdMPXV^Δ96,158^. Viral DNA from P10 was amplified by PCR, with pBAC-I^Δ96,158^ as a positive control. **F.** Read depth of P5 (red) and P10 (green) rdMPXV^Δ96,158^ aligned to the rdMPXV^Δ96,158^ reference genome. **G-H.** Read depth of P5 (red) and P10 (green) rdMPXV^Δ96,158^ aligned to MPXV reference genome, focusing on *OPG96* (G) and *OPG158* (H) and flanking regions.

**Supplementary Figure 4.**
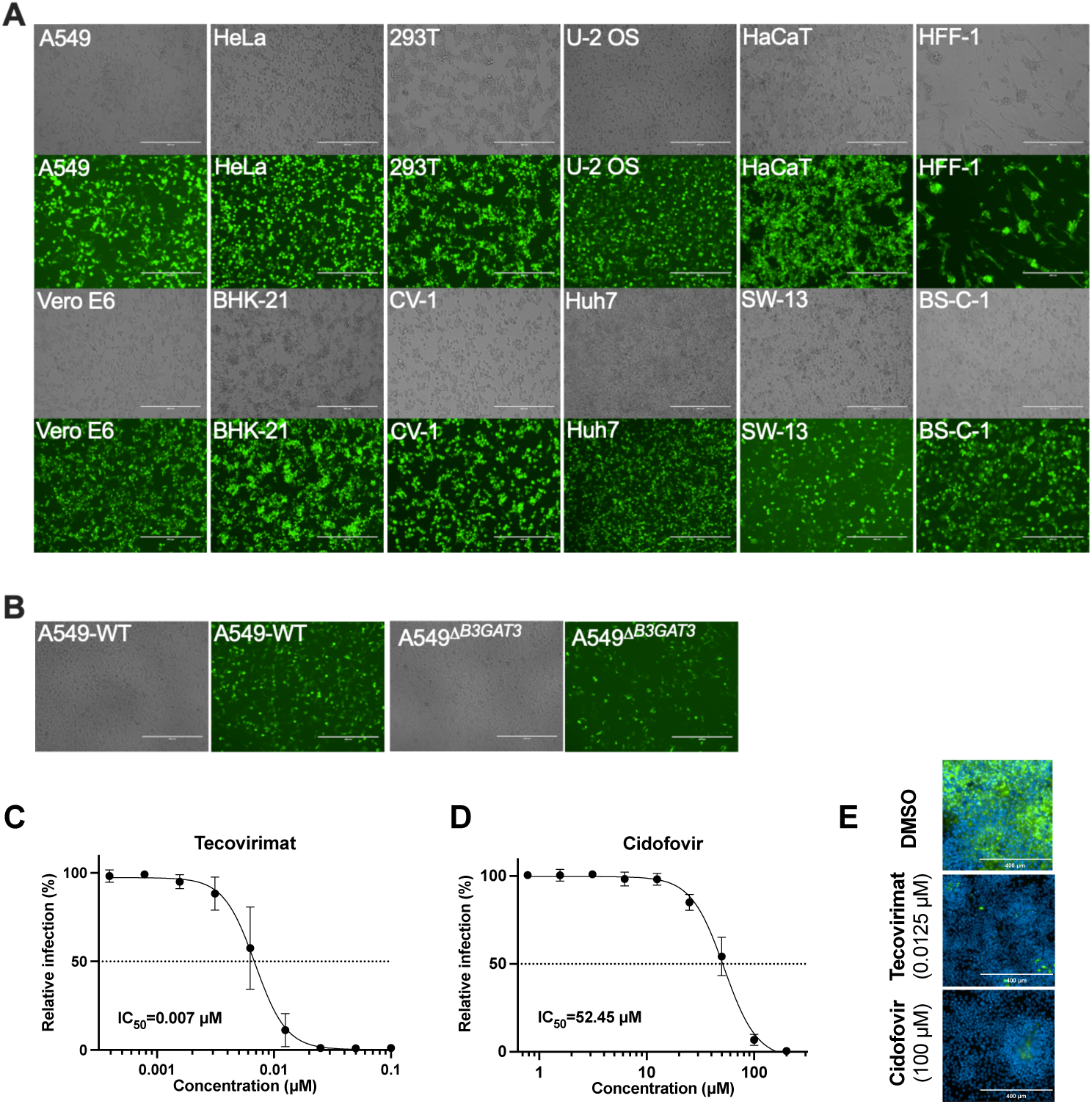
Cell susceptibility and antiviral compound assessment of replication-defective rdMPXV^Δ96,158^ particles. **A.** Representative images of rdMPXV^Δ96,158^ infection in various cell lines (MOI 0.5, 48 h), as described in Figure 2H). **B.** Representative images of rdMPXV^Δ96,158^ infection in WT and Δ*B3GAT3* A549 cells (MOI 0.5, 48 h), as described in Figure 2I. **C-D.** IC50 of tecovirimat (C) and cidofovir (D) in HaCat-96,158 cells infected with rdMPXV^Δ96,158^ (MOI 0.1, 30 h). **E.** Representative fluorescence images of HaCat-96,158 cells pre-treated with tecovirimat (0.125 μM) or cidofovir (100 μM) for 1 h, followed by rdMPXV^Δ96,158^ infection (MOI 0.1, 30 h). Cells were fixed, stained with DAPI, and imaged under a fluorescence microscope. Scale bar, 400 μm.

## REFERENCE

1. Relich RF, Loeffelholz MJ. Taxonomic Changes for Human Viruses, 2020 to 2022. J Clin Microbiol 61, e0033722 (2023).

2. Bunge EM, et al. The changing epidemiology of human monkeypox-A potential threat? A systematic review. PLoS Negl Trop Dis 16, e0010141 (2022).

3. Dimitrakoff J. Monkeypox Virus Infection across 16 Countries - April-June 2022. N Engl J Med 387, e69 (2022).

4. Van Dijck C, et al. Emergence of mpox in the post-smallpox era-a narrative review on mpox epidemiology. Clin Microbiol Infect 29, 1487–1492 (2023).

5. Isidro J, et al. Phylogenomic characterization and signs of microevolution in the 2022 multi-country outbreak of monkeypox virus. Nat Med 28, 1569–1572 (2022).

6. Otu A, Ebenso B, Walley J, Barceló JM, Ochu CL. Global human monkeypox outbreak: atypical presentation demanding urgent public health action. Lancet Microbe 3, e554–e555 (2022).

7. Du M, Liu M, Niu B, Liu J. The global alarm bell is ringing due to the threat of potential severe cases and deaths caused by clade I of monkeypox virus. Lancet Infect Dis, (2024).

8. Ndembi N, et al. Evolving Epidemiology of Mpox in Africa in 2024. N Engl J Med 392, 666–676 (2025).

9. Gessain A, Nakoune E, Yazdanpanah Y. Monkeypox. N Engl J Med 387, 1783–1793 (2022).

10. Senkevich TG, Yutin N, Wolf YI, Koonin EV, Moss B. Ancient Gene Capture and Recent Gene Loss Shape the Evolution of Orthopoxvirus-Host Interaction Genes. mBio 12, e0149521 (2021).

11. Vernuccio R, et al. Structural insights into tecovirimat antiviral activity and poxvirus resistance. Nat Microbiol 10, 734–748 (2025).

12. Yang G, et al. An orally bioavailable antipoxvirus compound (ST-246) inhibits extracellular virus formation and protects mice from lethal orthopoxvirus Challenge. J Virol 79, 13139–13149 (2005).

13. Wilkin T. Determining Effective Therapy for Mpox. N Engl J Med 392, 1547–1548 (2025).

14. Lenharo M. Hopes dashed for drug aimed at monkeypox virus spreading in Africa. Nature 632, 965 (2024).

15. Chan-Tack K, et al. Benefit-risk assessment for brincidofovir for the treatment of smallpox: U.S. Food and Drug Administration’s Evaluation. Antiviral Res 195, 105182 (2021).

16. Maruri-Avidal L, Weisberg AS, Bisht H, Moss B. Analysis of viral membranes formed in cells infected by a vaccinia virus L2-deletion mutant suggests their origin from the endoplasmic reticulum. J Virol 87, 1861–1871 (2013).

17. Maruri-Avidal L, Weisberg AS, Moss B. Direct formation of vaccinia virus membranes from the endoplasmic reticulum in the absence of the newly characterized L2-interacting protein A30.5. J Virol 87, 12313–12326 (2013).

18. Carten JD, Greseth M, Traktman P. Structure-Function Analysis of Two Interacting Vaccinia Proteins That Are Critical for Viral Morphogenesis: L2 and A30.5. J Virol 96, e0157721 (2022).

19. Smith GL, Vanderplasschen A, Law M. The formation and function of extracellular enveloped vaccinia virus. J Gen Virol 83, 2915–2931 (2002).

20. Li Q, Sun B, Chen J, Zhang Y, Jiang Y, Yang S. A modified pCas/pTargetF system for CRISPR-Cas9-assisted genome editing in Escherichia coli. Acta Biochim Biophys Sin (Shanghai) 53, 620–627 (2021).

21. Earl PL, Americo JL, Cotter CA, Moss B. Comparative live bioluminescence imaging of monkeypox virus dissemination in a wild-derived inbred mouse (Mus musculus castaneus) and outbred African dormouse (Graphiurus kelleni). Virology 475, 150–158 (2015).

22. Song G, Cheng L, Liu J, Zhou Y, Zhang C, Zong Y. Establishment of an animal model for monkeypox virus infection in dormice. Sci Rep 15, 4044 (2025).

23. Chung CS, Hsiao JC, Chang YS, Chang W. A27L protein mediates vaccinia virus interaction with cell surface heparan sulfate. J Virol 72, 1577–1585 (1998).

24. Hsiao JC, Chung CS, Chang W. Vaccinia virus envelope D8L protein binds to cell surface chondroitin sulfate and mediates the adsorption of intracellular mature virions to cells. J Virol 73, 8750–8761 (1999).

25. Lin CL, Chung CS, Heine HG, Chang W. Vaccinia virus envelope H3L protein binds to cell surface heparan sulfate and is important for intracellular mature virion morphogenesis and virus infection in vitro and in vivo. J Virol 74, 3353–3365 (2000).

26. Pokorny L, et al. The vaccinia chondroitin sulfate binding protein drives host membrane curvature to facilitate fusion. EMBO Rep 25, 1310–1325 (2024).

27. Sammon D, et al. Molecular mechanism of decision-making in glycosaminoglycan biosynthesis. Nat Commun 14, 6425 (2023).

28. Wolffe EJ, Vijaya S, Moss B. A myristylated membrane protein encoded by the vaccinia virus L1R open reading frame is the target of potent neutralizing monoclonal antibodies. Virology 211, 53–63 (1995).

29. Bojkova D, et al. Drug Sensitivity of Currently Circulating Mpox Viruses. N Engl J Med 388, 279–281 (2023).

30. Holzer GW, Falkner FG. Construction of a vaccinia virus deficient in the essential DNA repair enzyme uracil DNA glycosylase by a complementing cell line. J Virol 71, 4997–5002 (1997).

31. Mayrhofer J, et al. Nonreplicating vaccinia virus vectors expressing the H5 influenza virus hemagglutinin produced in modified Vero cells induce robust protection. J Virol 83, 5192–5203 (2009).

32. Boyle KA, Greseth MD, Traktman P. Genetic Confirmation that the H5 Protein Is Required for Vaccinia Virus DNA Replication. J Virol 89, 6312–6327 (2015).

33. Greseth MD, Czarnecki MW, Bluma MS, Traktman P. Isolation and Characterization of vΔI3 Confirm that Vaccinia Virus SSB Plays an Essential Role in Viral Replication. J Virol 92, (2018).

34. Warren RD, Cotter CA, Moss B. Reverse genetics analysis of poxvirus intermediate transcription factors. J Virol 86, 9514–9519 (2012).

35. Meng X, Wu X, Yan B, Deng J, Xiang Y. Analysis of the role of vaccinia virus H7 in virion membrane biogenesis with an H7-deletion mutant. J Virol 87, 8247–8253 (2013).

36. Hyun SI, Weisberg A, Moss B. Deletion of the Vaccinia Virus I2 Protein Interrupts Virion Morphogenesis, Leading to Retention of the Scaffold Protein and Mislocalization of Membrane-Associated Entry Proteins. J Virol 91, (2017).

37. Noyce RS, Lederman S, Evans DH. Construction of an infectious horsepox virus vaccine from chemically synthesized DNA fragments. PLoS One 13, e0188453 (2018).

38. Domi A, Moss B. Cloning the vaccinia virus genome as a bacterial artificial chromosome in Escherichia coli and recovery of infectious virus in mammalian cells. Proc Natl Acad Sci U S A 99, 12415–12420 (2002).

39. Wyatt LS, Earl PL, Moss B. Generation of Recombinant Vaccinia Viruses. Curr Protoc Protein Sci 89, 5.13.11–15.13.18 (2017).

40. Domi A, Moss B. Engineering of a vaccinia virus bacterial artificial chromosome in Escherichia coli by bacteriophage lambda-based recombination. Nat Methods 2, 95–97 (2005).

41. Tischer BK, Kaufer BB. Viral bacterial artificial chromosomes: generation, mutagenesis, and removal of mini-F sequences. J Biomed Biotechnol 2012, 472537 (2012).

42. Hao M, Tang J, Ge S, Li T, Xia N. Bacterial-Artificial-Chromosome-Based Genome Editing Methods and the Applications in Herpesvirus Research. Microorganisms 11, (2023).

43. Gietz RD, Schiestl RH. High-efficiency yeast transformation using the LiAc/SS carrier DNA/PEG method. Nat Protoc 2, 31–34 (2007).

44. Li H. Aligning sequence reads, clone sequences and assembly contigs with BWA-MEM. arXiv: Genomics, (2013).

45. Khanna A, et al. Bam-readcount - rapid generation of basepair-resolution sequence metrics. ArXiv, (2021).

